# Single Cell Spatial Chromatin Analysis of Fixed Immunocytochemically Identified Neuronal Cells

**DOI:** 10.1101/780387

**Authors:** Jaehee Lee, Youtao Lu, Jinchun Wang, Jifen Li, Stephen A. Fisher, C. Erik Nordgren, Jean G. Rosario, Stewart A. Anderson, Alexandra V. Ulyanova, Steven Brem, H. Isaac Chen, John A. Wolf, M. Sean Grady, Mimi Healy, Junhyong Kim, James Eberwine

## Abstract

Assays examining the open-chromatin landscape in single cells require isolation of the nucleus, resulting in the loss of spatial/microenvironment information. Here we describe CHEX-seq (**CH**romatin **EX**posed) for identifying single-stranded open-chromatin DNA regions in paraformaldehyde-fixed single cells. CHEX-seq uses light-activated DNA probes that binds to single-stranded DNA in open chromatin. *In situ* laser activation of the annealed probes’ 3’-Lightning Terminator™ in selected cells permits the probe to act as a primer for *in situ* enzymatic copying of single-stranded DNA that is then sequenced. CHEX-seq is benchmarked with human K562 cells and its utility is demonstrated in dispersed primary mouse and human brain cells, and immunostained cells in mouse brain sections. Further, CHEX-seq queries the openness of mitochondrial DNA in single cells. Evaluation of an individual cell’s chromatin landscape in its tissue context enables “spatial chromatin analysis”.

**One Sentence Summary:** A new method, CHEX-seq (**CH**romatin **eX**posed), identifies the open-chromatin landscape in single fixed cells thereby allowing spatial chromatin analysis of selected cells in complex cellular environments.

## Main Text

The process of RNA transcription requires a cell’s genomic DNA to be in an open-chromatin conformation, where there is less nucleosome packing, so that the transcription regulatory proteins can bind and function. It is clear that chromatin structure is dynamic and regulated by a number of factors including development, stress, and pharmacological challenge (*1–3*). Most chromatin modeling studies have relied upon the use of pooled cells to generate genomic DNA/chromatin for analysis. Included among chromatin analysis procedures are DNase-seq, FAIRE-seq, and ChIP-seq as well as other approaches. Recently, these methods have been extended to single cells (*4–7*). For example, the recent ATAC-seq approach to mapping chromatin in single cells exploits an assay for detecting transposon-accessible chromatin (*8*). This methodology uses Tn5 transposase to tag and purify accessible nucleosome-free double-stranded DNA regions in the genome. Each of these procedures has specific advantages and disadvantages, with the most significant disadvantage being that they all assess chromatin in nuclei isolated from the tissue of interest, thereby losing spatial location information and the cellular microenvironment context. To overcome this issue, we developed CHEX-seq (**<u>CH</u>**romatin **<u>EX</u>**posed) to assess chromatin conformation in fixed single cells including neurons and astrocytes.

### CHEX-seq Analysis

Open chromatin is composed of both double- and single-stranded DNA (*9–11*). The open state of chromatin is necessary for many cell functions, such as replication, homologous recombination, and DNA repair as well as transcription. However, “openness” may not correlate directly with transcription, as other trans-acting factors are also required (*12*). Single-stranded DNA is necessary for transcription in the form of the single-stranded “transcription bubble” which has been reported to be as large as ∼200 bases (*13, 14*). Further, in concert with transcription bubbles, transcriptionally active chromatin contains long stretches of single-stranded areas greater than a kilobase in length (*11, 14, 15*). The amount of single-stranded DNA in the genome is estimated to vary from ∼0.2% to 2.5%, depending upon the physiological state of the cell (*15*).

To assay single-stranded DNA at single-cell resolution *in situ*, we designed an oligonucleotide (Fig 1) that can anneal randomly to single-stranded genomic DNA and remain inactive until light-activation. After photoactivation, the oligonucleotide serves as a primer for DNA polymerase-mediated complementary DNA synthesis (Fig 1). This is DNA-directed *in situ* transcription (*16, 17*). The resolution of primer activation is determined by the diffraction limit of the activating light’s wavelength and the numerical aperture of the lens. To facilitate analysis, the CHEX-seq oligonucleotide was engineered to contain a sample-specific barcode: a T7 RNA polymerase promoter site along with a degenerate DNA sequence that is terminated with a fluorescently tagged, photo-reversibly blocked nucleotide (Fig 1A, Supplemental Data Fig 1). After DNA synthesis, the complementary DNA is removed with 0.1N NaOH, copied into double-stranded DNA, and linearly amplified using T7 RNA polymerase (aRNA amplification) (*18, 19*). The aRNA is subsequently reverse transcribed to 1^st^ and 2^nd^ strand DNA with custom primers, converted into a sequencing library, and sequenced (Supplemental Data Fig 1, Supplemental Data Fig 2).

**Figure 1.**
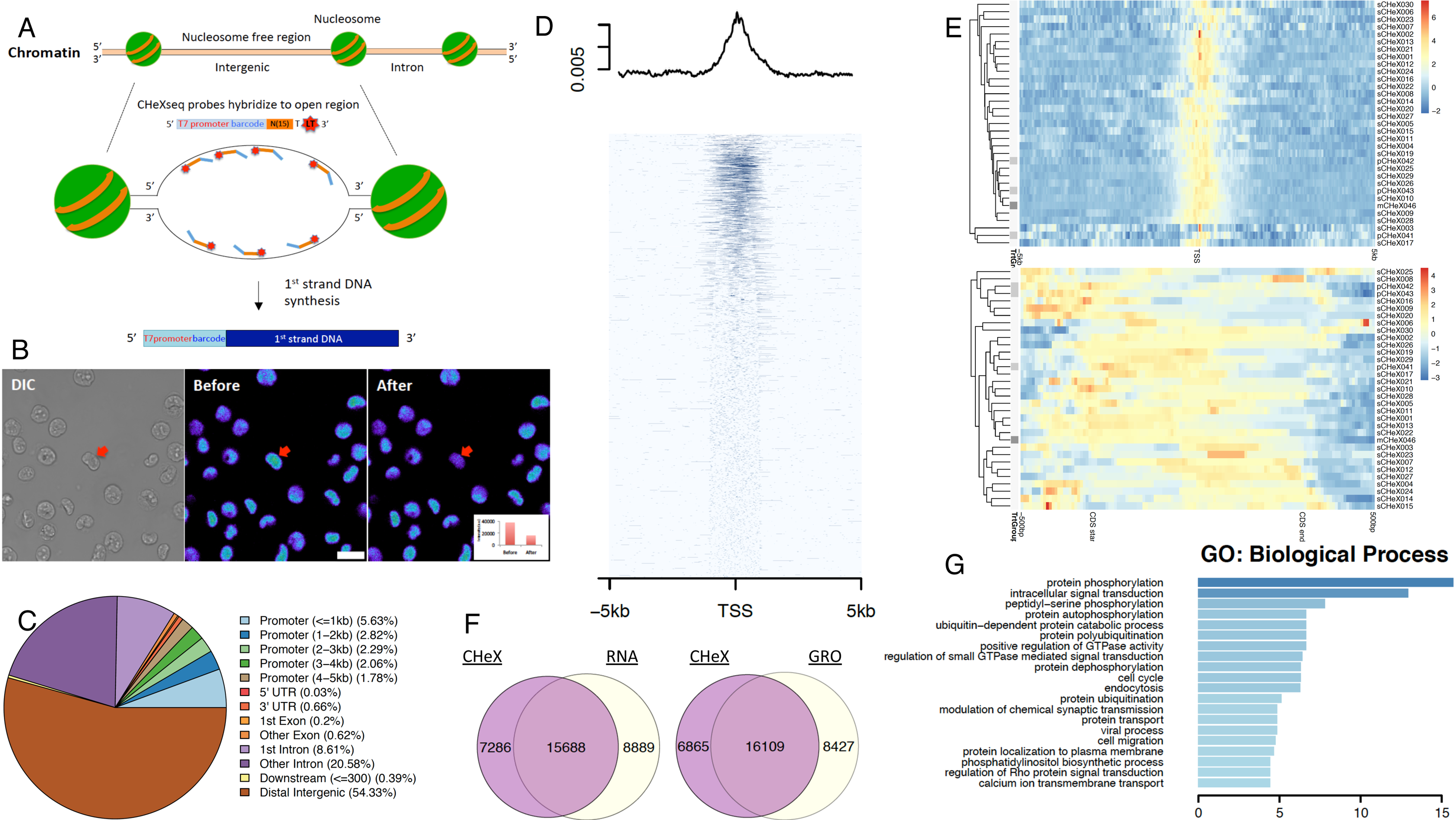
K562 CHEX-seq Benchmarking. (A) Schematic of CHEX-seq assay rationale, (B) CHEX-seq probe loading into K562 cell nuclei (DIC image) and fluorescence signal before and after activation of the CHEX-seq probe in a single nucleus (red arrow) scale bar= 20µm; (C) Statistics of CHEX-seq priming sites with respect to genomic features; (D) TSS proximal (+/-5kb) coverage of K562 samples (all positive samples merged); (E) z-scored coverage at TSS proximity (upper) and CDS (lower) at single-cell level; (F) Overlap between CHEX-seq primed genes (whole gene body > 0) and RNA-seq expressed genes (exon > median); (G) GO functional enrichment results (top 20) of the CHEX-RNA overlapping genes (subfigure F, left).

**Figure 2.**
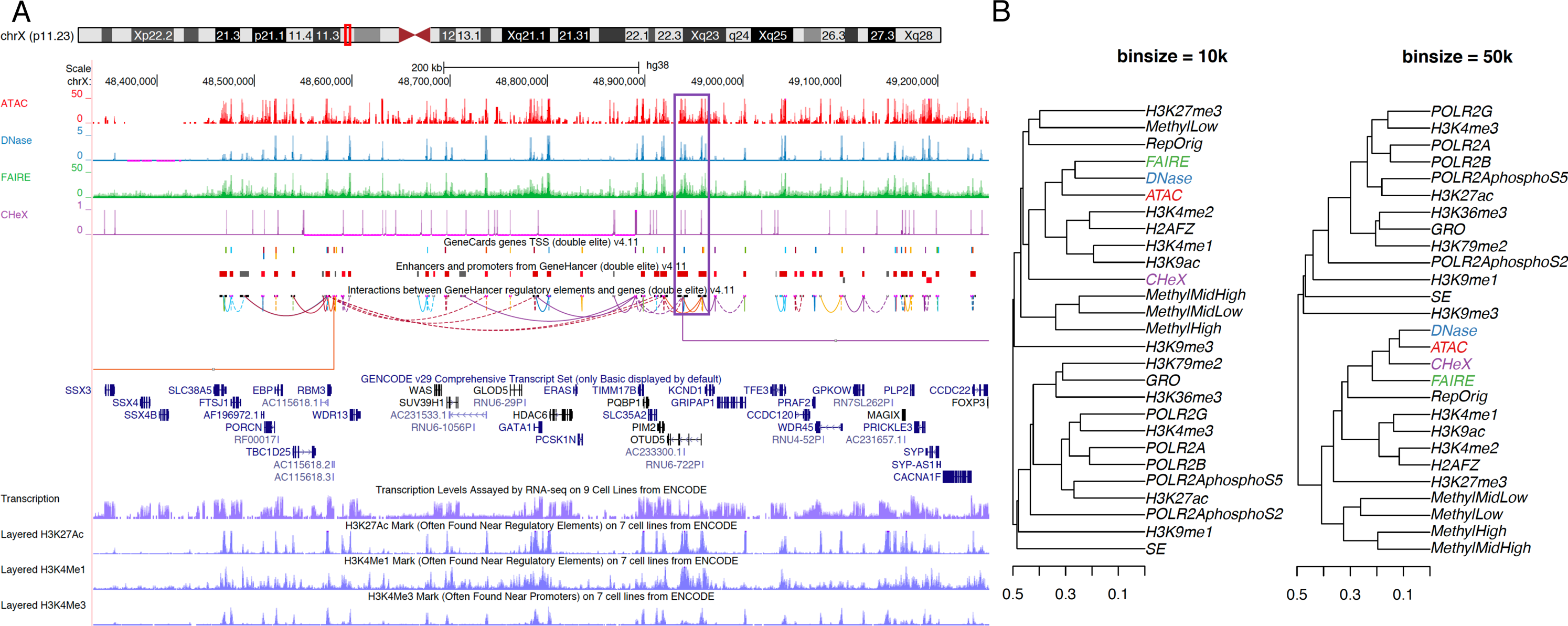
Genomic comparison of CHEX-seq with other open-chromatin assays. (A). UCSC Genome Browser track view comparing the coverage of CHEX-seq (purple) against ATAC-seq (red), DNase-seq (blue), FAIRE-seq (green) at locus OTUD5. Below the four assays are regulatory interaction tracks (GeneCards genes TSS, Enhancers and Promoters, and GeneHancer Proximal-Distal Interactions) derived from the GeneHancer database (*49*). The last four tracks are transcriptome and three histone marks (H3K27ac, H3K4me1, H3K4me3s). We note a regulatory interaction between OTUD5’s promoter and one of its 3’ introns is shared by all four open-chromatin assays (blue rectangle); (B). Hierarchical clustering of open-chromatin assays, transcriptome, and epigenomes at 10kb-bin (left) and 50kb-bin (right) resolution, using binarized coverage and Jaccard distance.

To show the utility of CHEX-seq we have chosen to assess CHEX-seq single-strand open-chromatin detection in multiple cell types including: two different species (human and mouse), the ENCODE K562 cell line, two types of neuronal cell preparations (dispersed neurons and *in situ* brain section neurons), two types of neurons (neocortical and hippocampal), two classes of brain cells (neurons and astrocytes) and finally cells of differing ages. This range of experimental samples highlights the general utility of the methodology and permits some interesting correlations to be examined.

### Benchmarking CHEX-seq in Human K562 cells

HK562 cells were selected for benchmarking the CHEX-seq procedure as this cell line was chosen by ENCODE for extensive analyses (The ENCODE Project Consortium, 2012). After fixation, K562 cells were gravity deposited onto poly-L-lysine-coated cover slips and then permeabilized and washed in PBS. Annealing of the CHEX-seq fluorescently labeled probes to the chemically fixed cells shows the probe concentrating in the nucleus of the cell (Fig 1B). The CHEX-seq primers were activated by illuminating with 405 nm (UV) laser at 60% power and 30µs per pixel, whereupon a 45∼80% decrease in fluorescence was observed (Fig 1B inset). This decrease is due to the loss of the fluorescent moiety and freeing of a 3’-hydroxyl group to prime DNA synthesis.

CHEX-seq reads were first preprocessed by a customized SCAP-T Next Generation Sequencing pipeline (https://github.com/safisher/ngs), then mapped back to the UCSC hg38 (human) or UCSC mm10 (mouse) genome. Finally, an additional QC procedure was applied to filter for good-quality reads (see Supplemental Text). We computed the percentage of CHEX-seq reads that map to gene features and the regions flanking genes, specifically including the 5’ promoter region, transcription start site (TSS), the 5’ untranslated region (UTR), exons, introns, the 3’ UTR, and the 3’-proximal area (Fig 1C). K562 cells show the highest proportion of CHEX-seq reads mapped to intergenic regions (>50%), followed by introns (∼30%) and then proximal promoters less then 1kb from the transcriptional start site (TSS) (∼6%). The promoter proximal region (< 1kb) of genes had 3 times more reads than distal regions (4-5kb), consistent with the opening of chromatin near the TSS. More specifically, TSS enrichment was observed in most single-cell samples, with weak or no enrichment in negative controls. Combining the signal across all non-control samples shows a distinct peak centered at the TSS (Fig 1D), much resembling the TSS peaks observed in ATAC-seq or DNase-seq assays (*8, 20*). ATAC-seq data shows a peak of sequencing reads around the TSS, while the CHEX-seq data has a similar peak with a slightly extended slope after the peak in the 5’ to 3’ direction. Fig 1E shows within-cell CHEX-seq signals from individual cells, pooled for annotated features. These data suggest a propensity for chromatin to be open near the start of the CDS (coding sequence). where there is an increased density of CHEX-reads.

To assess how many of the K562 CHEX-seq sites correspond to expressed mRNA, we compared the CHEX-seq data with published K562 transcriptome datasets (Fig 1F). These data showed that ∼64% of the expressed transcriptome (15,688 genes) had corresponding CHEX-seq sites. Even with this relatively large overlap, there were still 7,286 CHEX-seq genic regions (∼32% of CHEX-seq genes) that did not have evidence of transcription in public transcriptome data. In comparing CHEX-seq data to GRO-seq transcripts (a real-time transcription runoff assay (*21, 22*)), there was a similar number of overlapping genes (∼66%) and a decrease in the CHEX-seq unique genes. Since GRO-seq data is not dramatically influenced by RNA stability, it is a more accurate reflection of genes that are being actively transcribed from open-chromatin regions. Gene ontology analysis of CHEX-seq sites identified cell signaling, cell cycle, and GTPase regulatory pathways as enriched in K562 cells mRNA; consistent with the notion that K562 cells are a transformed cell line (Fig 1G). These data are consistent with the fact that K562 cells are a transformed cell line in which these pathways are functional.

The genome coverage of CHEX-seq data compared with other open-chromatin as well as epigenome assays is presented in UCSC Genome Browser (*23*) (Fig 2). As an example, the OTUD5 gene region shows the mapping of CHEX-seq reads compared to three open-chromatin assays (ATAC-, DNase-, FAIRE-seq), highlighting that each assay has both overlapping open-chromatin regions as well as regions unique to the method of analysis (Fig 2A). We note that there were 32 cells mapped for the CHEX-seq samples, versus ensemble mapping of more than 200 cells for the other assays. In this particular view of the gene, OTUD5, we note a regulatory interaction between OTUD5’s promoter and one of its 3’ introns that is shared by all four open chromatin assays (purple rectangle). Different epigenomic assays have different genomic scales due to both the biological nature of the signals detected by each technique and the different chemistry of the assays. To assess the relationship between different epigenomic assays, we computed signal concordance in two different window sizes for 27 different assays and clustered the results (Fig 2B). At the size scale of 10kb windows, the open-chromatin assays (FAIRE-, DNase-, and ATAC-seq) cluster together, while CHEX-seq is in the same cluster but at a larger distance; this cluster also includes histone methylation assays and the replication of origin assay. At a window size of 50kb, CHEX-seq, ATAC-seq, DNase-seq, assays form a cluster with FAIRE-seq just outside the cluster. As the average size of a human gene is ∼42kb and the functional transcriptional chromatin unit is ∼50kb (*24*), these data suggest that the same open-chromatin associated genes are identified with each of these procedures, but the single-stranded open-chromatin CHEX-seq positions are likely displaced from those of the other procedures. A direct overlap would not be expected, as the other procedures have a target bias for double-stranded DNA (ATAC-seq and FAIRE-seq) or are indiscriminate (DNase-seq) as compared to CHEX-seq’s single-stranded DNA requirement. The CHEX-seq signals may also be sparser due to limited numbers of analyzed cells, but these data highlight that both double-stranded and single-stranded DNA exist within the open-chromatin region, sculpting the open-chromatin landscape of a cell.

To confirm the single-stranded nature of a CHEX-seq predicted loci, we performed single molecule FISH for a CHEX-seq predicted K562 open-chromatin site on Chromosome 1 (630737–633960) (Supplemental Fig 3, Panel A). In addition to CHEX-seq identification of the open-chromatin status of this genic region, ATAC-seq predicts it to be open, while in contrast, DNase shows limited openness and FAIRE predicts that it is not open. Eight 20-mer oligonucleotide probes were synthesized to target this human Chromosome 1 area. These probes were labeled at the 5’-end with the ATT0 590 fluorophore. Upon performing FISH, generally 3 strong positive spots are observed in single cell nuclei (Supplemental Fig 3, Panel B). This trisomy signal is due to the complicated K562 cell karyotype where some cells have 3 copies of Chromosome 1 (*25*).

**Figure 3.**
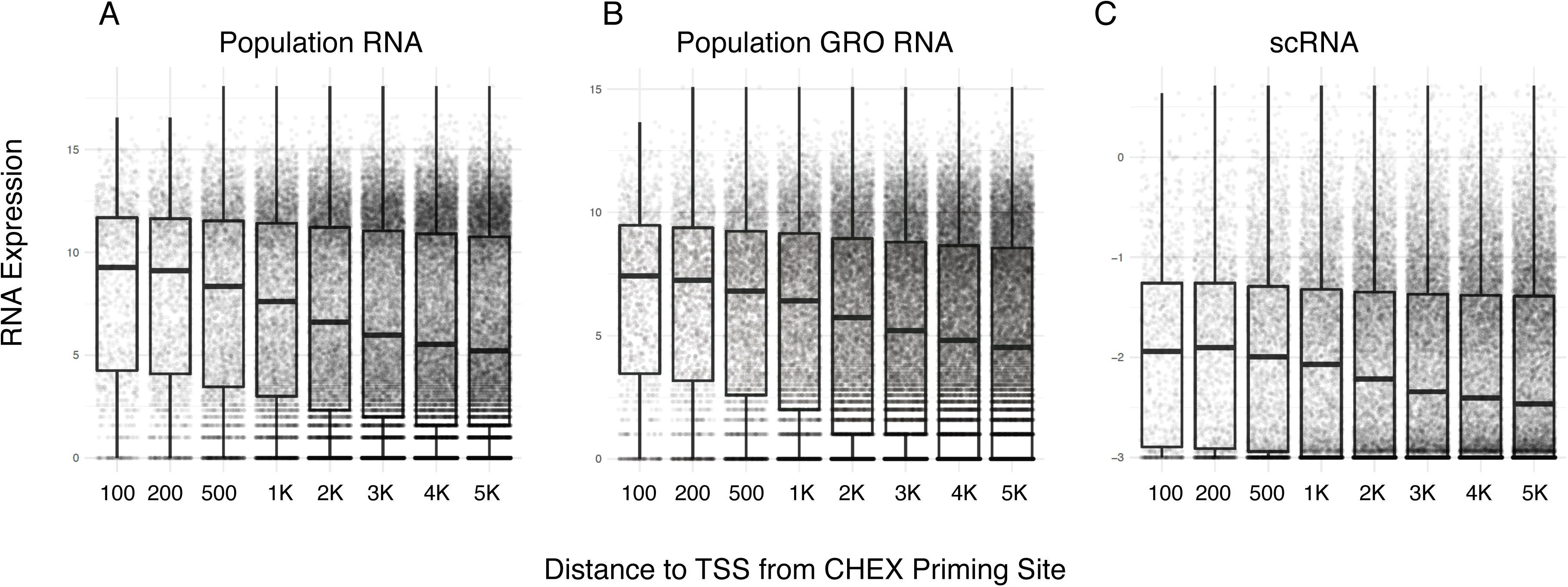
Correlation of CHEX-seq read distance from TSS with gene RNA abundance. (A). Bulk K562 RNA-seq; (B) Bulk K562 GRO-seq; (C) K562 scRNA-seq, single cells averaged. Y-axis: gene expressions; x-axis: distance to TSS from CHEX priming sites.

We next stratified K562 CHEX-seq priming sites with respect to their distance to the cognate genes’ TSS and compared this distance with mRNA expression level, from the same gene (Fig 3). Using three different mRNA sources – population mRNA (GSE32213), GRO-seq mRNA (GSE60454) and single-cell mRNA (scRNA, GSE90063) – it can be seen that when the CHEX-seq priming sites are closer to the TSS, the corresponding mRNAs are generally present in higher abundance (*26*). This pattern is found in human neurons and astrocytes, and mouse neurons, astrocytes, and section-localized neurons (Supplemental Data, Figure 4). These data suggest a regulated plasticity with regard to single-stranded DNA within a gene: i.e., as transcription moves along the length of the gene, the 5’-open site becomes unavailable for hybridization, perhaps due to reannealing of the single-stranded region. In this model, detectable CHEX-seq priming sites have varying half-lives, and those that are proximal to the TSS remain single-stranded for a longer time and correspond to high levels of transcription. Thus CHEX-seq priming closer to the TSS would be more predictive of highly transcribed genes. These data are not simply due to differences in the rate of RNA stability, as GRO-seq RNA detects newly synthesized nuclear RNA. One model is that the TSS proximal single-strandedness is associated with gene expression, whose accessibility decays with precession of transcription, while more distal regions’ single-stranded accessibility might be related to other conformational regulation of the DNA.

**Figure 4.**
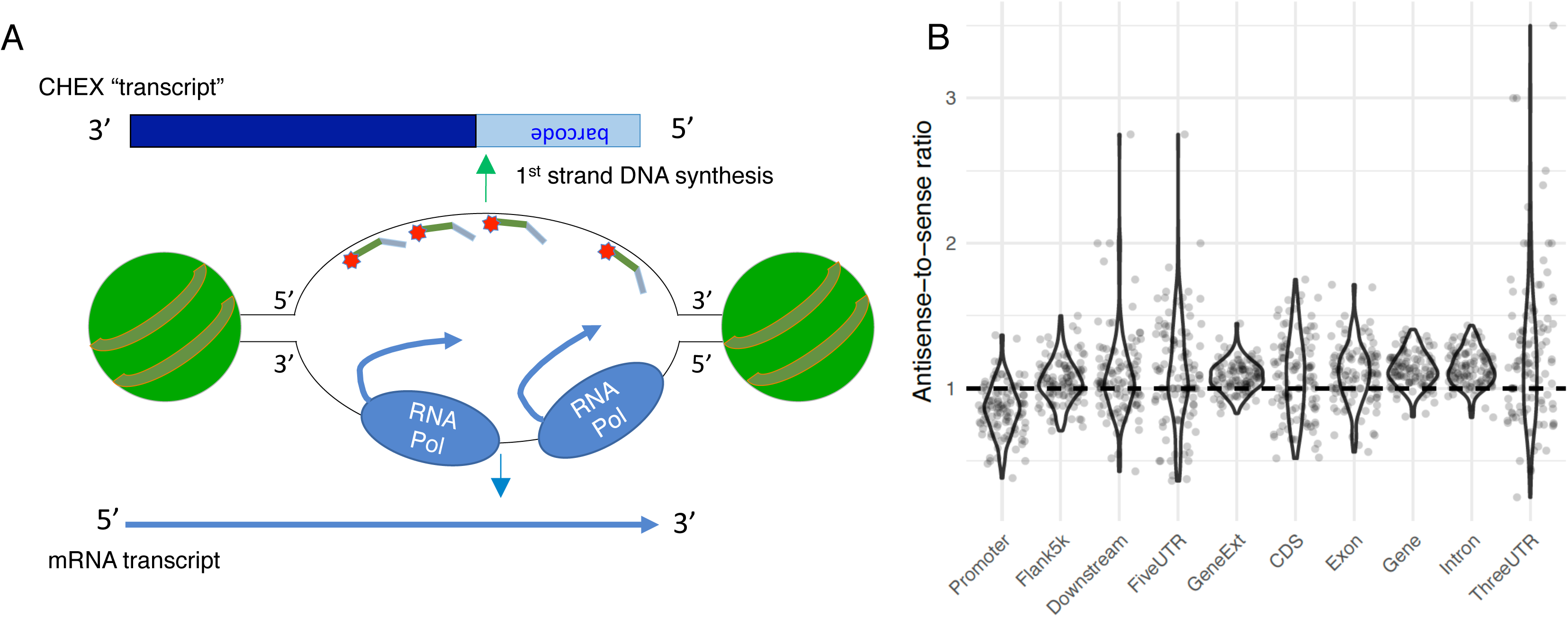
CHEX-seq Strandedness: detecting open chromatin’s strandedness. (A) Schematic showing the hypothesis that CHEX-seq priming-extending products should have opposite strandedness from sense-strand mRNA transcripts; (B) testing the hypothesis in (A). X-axis: various genomic features where CHEX-seq priming events are counted and binarized; y-axis: ratio of number of antisense- stranded over sense-stranded CHEX-seq products.

Since the RNA Pol2 transcriptional complex binds to the template DNA and synthesizes RNA transcripts in a 5’ to 3’ direction by transcribing the antisense strand, we wanted to assess whether or not CHEX-seq probes might be preferentially bound to the potentially more accessible sense strand, giving rise to an excess of “antisense-strand” reads (see schematic in Fig 4A). Fig 4B shows the ratio of antisense to sense reads for different annotated regions of the gene model. As expected, we found more antisense-stranded reads in the transcribed regions of the genome, with slightly increasing bias from the 5’ UTR towards the 3’ UTR. Interestingly, the promoter region exhibited an opposite bias toward sense-strand CHEX-seq reads (Fig 4B). This may be reflective of the antisense-strand being bound to proteins including Pol2 as it copies the antisense template, leaving the sense-strand more available for CHEX-seq primers to bind (*27, 28*). We speculate this opposite trend in promoters to be related to bidirectional promoter activity (*29*).

### *In Situ* Mouse Brain Tissue Section and Dispersed Single Neuron Analysis

To identify open-chromatin sites in individual neurons *in situ* in adult mouse brain tissue, where the neurons are in their natural context, we applied CHEX-seq to fixed adult brain tissue sections (100 µm) that were labeled by immunofluorescence with an antibody that detects neuronal microtubule associated protein 2 (MAP2). CHEX-seq probes were then annealed to the single-stranded DNA in the tissue section (for schematic see Fig 5A). Fig 5B shows the CA1 region of the hippocampus labeled for MAP2 immunofluorescence (green) and the CHEX-seq probe (red). The CHEX-seq probes were activated (confirmed by the loss of fluorescent signal) in an individual nucleus (arrow in boxed area of Fig 5B) after which *in situ* copying of DNA from single-stranded genomic DNA was performed. The CHEX-primed DNA was removed, amplified and sequenced. In comparing the open-chromatin CHEX-seq sites from *in situ section-localized* neurons with the expressed transcriptome from single cells (Fig 5D), there is a 59% overlap of CHEX-seq sites with expressed mRNA, while 88% of the transcriptome overlaps with CHEX-seq reads. This leaves 41% of CHEX-seq sites as not detected in mRNA while only 13% of the transcriptome does not show CHEX-seq open-chromatin sites. These data show that there is a large amount of single-stranded open-chromatin in fixed tissue sections that is not represented in the transcribed mRNA, likely corresponding to genes that are ready to be transcribed, non-messenger RNA, DNA replication sites or other types of DNA organizational structures. CHEX-seq reads in the tissue section can be further broken down to show an overlap with the transcriptome of 69% for exonic regions and 65% for intronic regions. This overlap suggests that the chromatin landscape and transcriptome are well correlated in cells that are localized in their natural microenvironment.

**Figure 5.**
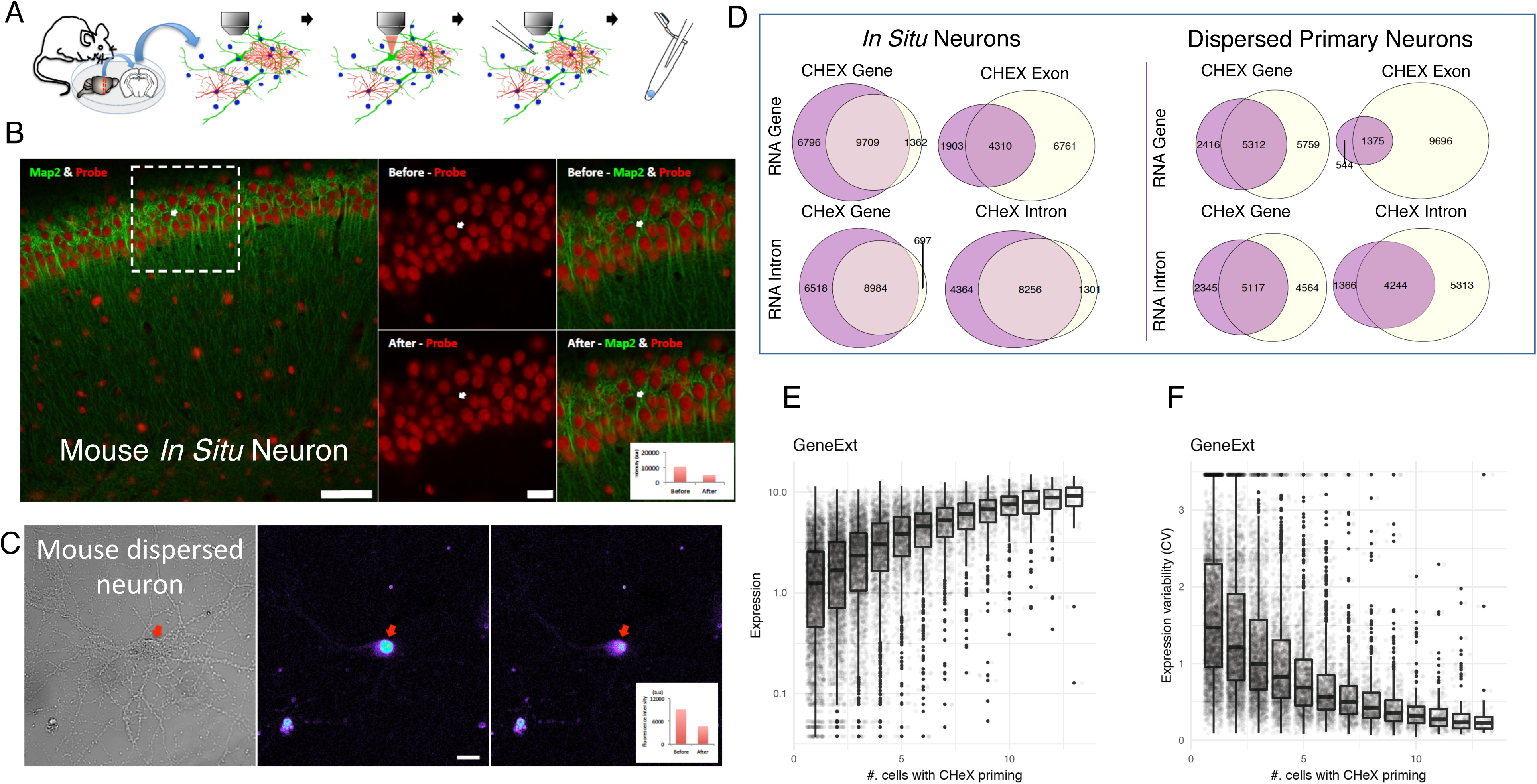
CHEX-seq analysis of single neurons in fixed mouse tissue sections and dispersed cell culture. (A) Schematic of CHEX-seq analysis of paraformaldehyde-fixed tissue sections. (B) Hippocampal section showing neurons immunolabeled for MAP2 (green). Red fluorescence indicates localization of the CHEX-seq probe. The right-most panels show reduced fluorescent signal in the single neuronal nucleus (white arrow) that was activated. scale bar= 20µm. (C) Paraformaldehyde-fixed, cultured cortical neuron, shown by DIC microscopy (left panel) and nuclear fluorescence for the CHEX-seq primer (middle panel). This signal is diminished after probe activation (right panel; quantified in the right panel insert). scale bar= 20µm. (D) CHEX-transcriptome comparison. Left, mouse fixed tissue section; right, mouse dispersed neurons. Rows are scRNA-seq average expression in exonic or intronic region, columns are CHEX-seq binarized priming signal in whole gene body, exonic or intronic region. (E) Correlation in intronic regions between CHEX-seq priming frequency and transcriptional <u>activity</u> in hippocampal sections. (F) Correlation in intronic regions between CHEX-seq priming frequency and transcriptional <u>variability</u> in mouse slice tissue.

In order to assess the pattern of single-stranded open-chromatin regions in dispersed neuronal cells, we also examined single fixed cultured mouse neurons (Fig 5C). As adult mouse neurons cannot be cultured and hippocampal cells are difficult to culture, we assessed open-chromatin sites in mouse neonate cortical neurons that were in primary culture for two weeks, during which time they develop dendrites. The dispersed mouse cortical neurons displayed TSS peaks similar to those observed for K562 cells (Fig 1D) as well as other cell types (Supplemental Data, Fig 5), which exhibit the same TSS open-chromatin conformation. However, we found fewer total CHEX-seq reads mapping to the expressed transcriptome in the dispersed cortical neurons (5, 312) as compared with the *in situ* hippocampal neurons (9,709) (Fig 5D). We found 88% of the transcriptome mapping onto CHEX-seq reads for the *in situ* neurons and only 48% for the dispersed neuron transcriptome. However, the percent of CHEX-seq reads that correspond to transcribed mRNA is 68% for dispersed cells as compared with 59% for *in situ* neurons. In general, a higher percentage of CHEX-seq positive regions show evidence of transcription in dispersed cortical neurons as compared to *in situ* hippocampal cells, while a markedly lower percentage of transcribed mRNA show CHEX-seq evidence in dispersed culture compared to section. While it is difficult to discern the relative contribution of cell type (although neocortical and hippocampal cell transcriptomes are very similar(*30*)) and cell age to these data, one potential interpretation is that there are more non-transcription associated open-chromatin sites in brain section neurons than in dispersed neurons.

In comparing the mouse dispersed cortical neuron CHEX-seq data with the averaged transcriptome of single cortical neurons there is a large overlap of the single-stranded DNA sites with transcribed mRNA (Figure 5D, right panel, upper-left). Of the 7,728 CHEX-seq active sites, 69% overlap with the transcriptome, leaving 31% of the single-stranded sites showing no detectable transcribed mRNA. Concomitant with these data of the 11,071 different transcripts identified in the single cells, 48% correlate with single-strand open-chromatin genes. To investigate the systems aspect of this comparison, we looked at Gene Ontology (GO) enrichment in genes in common as well as unique to either assay in dispersed neurons. There are 235 GO Molecular Function terms enriched at a Benjamini-Hochberg (BH) adjusted p-value of <0.1, shared between the open-chromatin analysis and transcriptome, while at the same significance, there are 40 in the CHEX-seq unique genes and 107 in the transcriptome unique genes. Among the shared pathways are those for *chromatin binding* (p-value: 2.0×10^-14^), *calmodulin binding* (p-value: 1.9×10^-10^) and many associated with neuronal function.

Evidence for the enrichment of these pathways in both the open chromatin and transcriptome of neurons is not surprising, as they give rise to normal cellular function as well as some of the specialized functions of neurons.

Interestingly, for *in situ* neurons, there was a significant relationship between the expression level of RNA and the number of CHEX-seq priming sites within that gene (Fig 5E). These data suggest that the number of CHEX-seq priming sites in a gene is indicative of the amount of transcription from that gene, with more sites suggesting more transcription. This relationship was somewhat surprising, as steady state levels of RNA are dependent upon other factors in addition to transcriptional activity, such as RNA stability. A single open site can correlate with high levels of expression (Fig 5E left panel, left-most) but such sites are much fewer in number. The large number of open-chromatin single-stranded sites in highly expressed genes of cells in the tissue section may be reflective of a higher level of activity where a gene is bursting transcriptional activity more frequently and the open-chromatin single-stranded state is maintained for an extended period of time. This is consistent with data showing that the variability in gene expression decreases when there are more CHEX-seq priming sites (Fig 5F). These data suggest that mean-scaled variability of expression may be inversely related to the quantitative degree and base-pair span of single-stranded DNA regions. Thus, the CHEX-seq priming measure may correlate with temporal constancy of transcription as well as overall production levels, which would be reflective of the cells’ higher metabolic needs and requirement for constant high levels of expressed RNAs.

We examined priming rates in units of extended genic regions defined as the whole transcribed region (5’UTR, exons, introns, 3’UTR) with an additional 5kb both upstream and downstream. For each extended genic region, we pooled the priming events from 28 cultured neuronal samples and 15 *in situ* hippocampal neuronal samples and carried out Fisher’s exact test for differential proportions, given the total reads in each treatment. We found a total of 86 significantly different priming rates (i.e., single-stranded regions) in extended gene regions after multiple test correction (Benjamini-Hochberg adjusted p-value < 0.05); there were 50 genic regions with greater CHEX-seq priming rate for dispersed cortical neurons versus *in situ* hippocampal neurons and 36 genic regions with greater priming rate *in situ* compared to dispersed culture.

The 50 genic regions with greater priming rates for cortical neurons in dispersed culture included a diverse set of gene functions. It appears that there is a shift in biology upon dispersion, with dispersed cell genes showing more single-strandedness for GO-annotated genes associated with cilium function, membrane function and nucleotide binding. Since many genes in these functional classes are involved in cell shape in yeast (*31*), these data suggest that upon cell dispersion, shape-altering genes might be activated. When we examine these 50 genic regions for that correspondence with the single cell transcriptome from dispersed cells, two of the genes that showed higher read recovery in dispersed cells were ACOX3 (Acyl-coenzyme A oxidase 3, an enzyme that functions in the peroxisome (*32*)) and SUDS3 (a subunit of HDAC1-dependent SIN3A co-repressor complex (*33*)). SUDS3 is thought to repress transcription by augmenting HDAC1 activity through modulation of chromatin structure. It is possible that SUDS3 protein is increased in dispersed cells and would function to decrease the number of open-chromatin sites upon dispersion. This may be especially true for the large number of non-transcribed CHEX-seq accessible single-stranded open-chromatin sites identified in section hippocampal neurons.

The 36 genic regions with greater priming rate in *in situ* hippocampal cells were concentrated on mitochondria-encoded genes, with 27 out of the 37 mitochondria-encoded genes showing significant differences (Supplemental Data Table 2). Mitochondrial DNA has been noted in other open-chromatin assays but has generally been removed for nuclear DNA analysis (*34*). We note that mitochondrial DNA is not organized into chromatin, as nuclear DNA is, but rather has a nucleoid structure (containing single-stranded DNA regions) that is dynamically regulated and transcribed (*35–37*). For these genic regions, the neurons from the fixed section showed an average of 15.7 CHEX-seq priming events per gene per cell, ranging from 6.8 events/cell to 32.1 events/cell. Compared to these values, we found only 0.016 average priming events per cell per mitochondrial-encoded gene for neurons from culture. Since CHEX-seq priming is limited by the interval of single-stranded regions, we do not expect a very large number of priming events per genic region in general, and we hypothesize these events to represent the single-stranded DNA found in multiple mitochondria in a given cell. These results indicate that mitochondrial activity, mitochondria replication and/or gene transcription, may be reduced in neurons in culture. We note that there were four cells (single-cell samples) in tissue sections that also had almost no CHEX-seq priming in these 27 mitochondrial gene regions, while showing strong signal from other genic regions, suggesting that the mitochondrial DNA activity states are heterogeneous between individual cells.

### Mouse and Human Astrocytes Promoter Openness

We further performed the assay on neonatal mouse and adult human astrocytes that were in culture for two weeks to compare against neurons of the same species and age (Supplemental Data Fig 6A). Mapping the CHEX-seq reads to the annotated gene model, astrocytes have a higher proportion of CHEX-seq reads in the promoter region of genes than neurons from the same species (Supplemental Data Fig 6B). These data are in accord with earlier studies (*38*) showing that the chromatin landscape of dividing cells (astrocytes) has more DNase I sensitive open chromatin around the promoter region of genes than that of terminally differentiated cells (neurons). This is particularly intriguing as the cells cross a wide age span, with the mouse cells being neonatal and the human cells originating from subjects ranging in age from 50-70 yrs. We noted above that the promoter-proximal CHEX-seq priming is more indicative of gene transcription.

**Figure 6.**
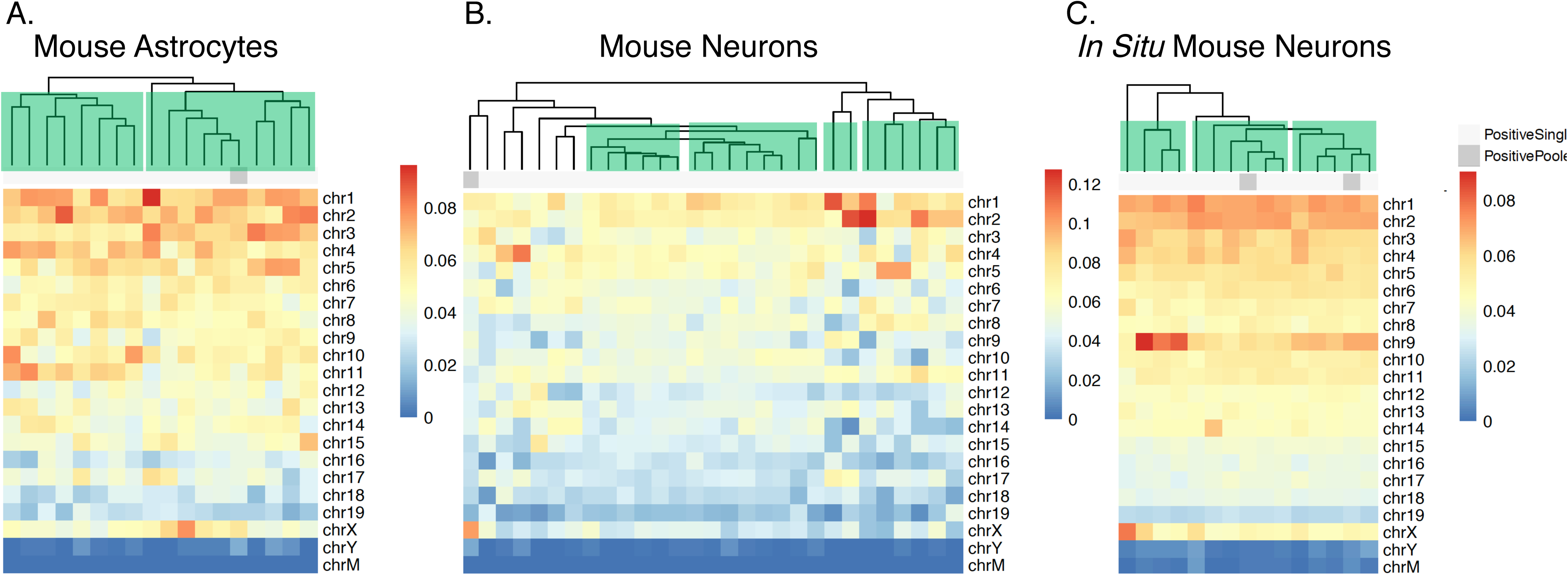
Chromosomal Landscape of Single-Stranded Open-Chromatin Between Cell Types. Distribution of CHEX priming sites by chromosome; color: fraction of priming frequency per chromosome. (A) mouse astrocyte culture, (B) mouse dispersed neuron cells, (C) mouse neuron section.

### Open-Chromatin Landscape Across the Mouse Genome

As CHEX-seq provides a whole genome view of single-stranded open-chromatin, we tested whether there was differential chromosome accessibility between mouse neurons and astrocytes. In Figure 6 the CHEX-seq read density for all of the chromosomes (rows) for each of the individual cells (columns) is plotted as a heatmap. Two things come to the fore in looking at these data: 1) the different cell types show different single stranded open-chromatin densities across the chromosomes; and 2) within a cell type there are groupings of cells that likely correspond to subtypes of the parent cell type. The *in situ* localized hippocampal neurons (Fig 6, far right panel) have a greater density of reads on chromosomes 1, 2 and 9 then the dispersed cortical neurons or astrocytes. Further, there are three subgroupings of these *in situ* neurons, with one group showing less chromosome 9 read density (green rectangles highlighting dendrogram groupings). The astrocytes likewise can be grouped into at least three groups (Fig 6, far left panel) with one of the discriminators being the density of open-chromatin on chromosome 11. As there are cells from multiple animals in each of the groupings, the groupings are not due to batch effects. Indeed, these data highlight the ability to characterize cell types based upon open-chromatin status. Why the chromosomal open-chromatin landscape exhibits differences between cells is unclear, but these data reflect the dynamism of the genome. Future studies will elicit a more finely detailed map of single stranded open-chromatin DNA dynamics.

### Summary and Future Possibilities

CHEX-seq queries single-stranded open-chromatin DNA regions and thus is distinct from but complementary to other chromatin analysis procedures, such as ATAC-seq that queries double-stranded DNA, and DNase I analysis that cuts both single and double-stranded DNA. As an interrogator of single-stranded DNA, CHEX-seq not only assesses nuclear DNA but also queries the single-stranded open-chromatin status of the mitochondrial genome. Surprisingly, there is strandedness availability of single-stranded genomic DNA to bind to complementary primers, suggesting a scaffolding of proteins and DNA that likely plays a role in the regulated transcription of DNA. Importantly, there was a high degree of overlap of CHEX-seq detectable open-chromatin sites and proximity of CHEX-seq priming sites with the expressed transcriptome, which may be useful in reporting the interrelated regulation of chromatin dynamics and transcription. CHEX-seq also provides evidence for genomic DNA regions that exhibit single-strandedness but are not transcribed, potentially including areas of DNA repair and sites of replication in dividing cells (*12, 39*). The potential for CHEX-seq to query these other genomic DNA sites awaits future studies. The ability to perform single cell open-chromatin analysis on immuno-identified spatially-localized neuronal cells (i.e. “Spatial Genomics”) promises to provide high resolution genomic analysis of dynamic circuit function and disease-associated dysfunction in functionally relevant cells.

As it is possible to synthesize CHEX-seq probes with different barcodes, it should be possible to multiplex these probes to transform the CHEX-seq technique into a moderate/high throughput methodology providing spatially defined chromatin landscape information for 10,000’s of cells from the same brain tissue section. Further, with selected enzymes and appropriate experimental conditions, it should also be possible to combine genomic DNA chromatin analysis with the transcriptome in the same cell. Finally, the ability of CHEX-seq to work with chemically fixed tissue presents the possibility of analyzing the open-chromatin landscape of individual cells or ensembles of cells in archival human post-mortem fixed tissues from various control and disease states. This suggests the intriguing possibility that preserved DNA-containing ancient specimens may retain some of their single-stranded chromatin structure (due to preservation), thereby providing the potential for CHEX-seq to define the open-chromatin status of these samples and provide insight into the expressed transcriptome from these ancient cells.

## Acknowledgements

We thank Eun-Hee Shim for technical help in the early stages of this work and Kevin Miyashiro for culturing the mouse neuronal cells.

## Funding

This work was funded in part by NIH U01MH098953 (J.E., J.K.) and RM1HG010023 (J.K., J.E.), and by R01MH110185 (S.A).

## Authors contributions

J. Lee, J. Li, J.R. conceived of and performed experiments; Y.L., S.F., C.E.N., J.Y.K. conceived of and performed data analysis; J.W., M.H., J.E. conceived of and/or performed CHEX primer synthesis; A.U., S.B., H.C., J.W., M.G. provided human neurosurgical tissue; S.A. provided immunostained brain sections; J.Y.K., J.E. conceived of experimental paradigm; J.Lee, Y.L., J.W., J. Li., S.F., C.E.N., J.R., S.A., A.U., S.B., H.I.C. J.W., M.S.G., M.H. J.K. J.E. wrote and reviewed the manuscript.

## Competing interest

J.E., J.Y.K., Y.L., S.F., J. Li,, J.W. and M.H. are co-inventors on a submitted CHEX-seq patent.

## Data and materials availability

Data will be deposited into public databases upon acceptance of the manuscript.

## Supplemental Materials

### Material and Methods

#### Human Brain Tissue

Human brain tissue was collected at the Hospital of the University of Pennsylvania (IRB#816223) using standard operating procedures for enrollment and consent of patients. Briefly, an en bloc sample of brain (typically 5×5×5 mm) was obtained from cortex that was resected as part of neurosurgical procedures for the treatment of epilepsy or brain tumors. This tissue was immediately transferred to a container with ice-cold oxygenated artificial CSF (in mM: KCl 3, NaH_2_PO_4_ 2.5, NaHCO_3_ 26, glucose 10, MgCl_2_-6H_2_O 1, CaCl_2_-2H_2_O 2, sucrose 202, with 5% CO_2_ and 95% O_2_ gas mixture) for transfer to the laboratory. Tissues arrived in the laboratory ∼10 minutes post excision. The brain tissues were then processed for cell culturing and fixation (see below).

#### Cell Culturing/Preparation and Fixation

K562 cells were obtained from ATCC and cultured in RPMI 1640 medium (Invitrogen) with 10% FBS and penicillin-streptomycin in a T75 flask at 37 °C in 5% CO2 for 2∼3 days. The cultured cells were transferred to a 50 ml tube and 16% paraformaldehyde (final 1%) was added for 10 mins at room temperature to fix the cells. After fixation, 1 M glycine (final 200 mM) with RPMI 1640 medium was used to quench for 10 mins followed by centrifugation at 300 x g for 5 mins. The supernatant was discarded and 3 mL of PBS were added to the pellet and then mixed by gently pipetting up and down 10-15 times using a fire-polished glass-pipette, to prevent cell clumping, and centrifuged at 300 rpm for 5 mins. The 100 µl cell pellet was attached to 18 mm gridded coverslips by incubating them for 2 hr. at room temperature. The samples were treated with PBS (w/o Ca^++^, Mg^++^) containing 0.01% Triton X-100 for 10 mins and then washed with PBS (w/o Ca^++^, Mg^++^) 3 times for 3 mins. To prepare human neuronal cell cultures, adult human brain tissue was placed in the papain (20 U, Worthington Biochemical) solution to dissociate at 37 °C for 30 to 40 mins and followed by ovomucoid (a papain inhibitor, 10 mg/ml, Worthington Biochemical) to stop the enzymatic dissociation (*40*). The tissue was triturated with a fire-polished glass Pasteur pipette. The cloudy cell suspension was carefully transferred to a new tube and centrifuged at 300 x g for 5 mins at room temperature. The cells were counted in an Autocounter (Invitrogen). Cells were plated on poly-L-lysine-coated (0.1 mg/ml, Sigma-Aldrich) 12-mm coverslips at a density of 3 × 10^4^ cells/coverslip. Cultures were incubated at 37 °C, 95% humidity, and 5% CO_2_ in neuronal basal medium (Neurobasal A, Gibco), serum-free supplement (B-27, Gibco) and 1% penicillin/streptomycin (Thermo-Fisher Scientific).

Dispersed mouse neuron/astrocyte cultures were prepared following published protocols (*41*). Dispersed cells were fixed using 4% paraformaldehyde for 10 min at room temperature. This was followed by three washes with 1x PBS. The cells were permeabolized with 0.1% Triton-X100 for 10 min at room temperature followed by three washes with 1x PBS.

#### Mouse brain tissue section preparation

A 3-month old male mouse was anaesthetized with halothane, euthanized by thoracotomy, then subjected to cardiac perfusion with 5 ml PBS followed by 20 ml PBS/4% paraformaldehyde. The brain was removed and post fixed at 4 °C for 16 hr., then rinsed in PBS and sectioned in the coronal plane at 100 µm on a vibratome (Leica VT-1000s). Sections including the hippocampus were then subjected to immunofluorescence labeling with chicken anti-MAP2 antisera (1:1000; Ab 5392; Abcam) followed by Alexa 488 conjugated goat anti-chicken secondary antibody (1:400; ab150169; Abcam).

#### CHEX-seq Probe Synthesis

HPLC-purified probe oligo and its complimentary oligo were purchased from Integrated DNA Technologies (IDT). A template-dependent DNA polymerase incorporation assay was employed to extend Cy5-dye-labeled Lightning Terminator™ (Agilent, Inc.) to the 3’ end of probe oligo: (1) 5 µM of probe oligo, 25uM complimentary oligo, 50 µM of Cy5-labeled Lightning Terminator™, 4 mM MgSO4, and 0.1U/µL of Therminator (New England Biolabs) were mixed in 1x ThermoPol buffer, (2) the mix was heated to 80°C for 45 sec and (3) then incubated for 5 mins at each of 60°C, 55°C, 50°C, 45°C, 40°C, 35°C, 30°C and 25°C. The incorporation product was purified on the 1260 Infinity reverse phase HPLC (Agilent Technologies) using the XTerra MS C18 Prep column (Waters). The purified product solution was concentrated to approximately 250 µL using the Vacufuge (Eppendorf) followed by denaturation into single-stranded oligo with equal volume of 0.2 M NaOH. HPLC purification and concentration were repeated using the same conditions for collection of the Lighting Terminator-labeled single-stranded probe. The final product was dissolved into 1xPBS and the concentration was determined by measuring Cy5 absorbance at 647 nm (Supplemental Data, Fig 1).

#### CHEX-seq probe application

After fixation and permeabilization, the cells and brain slices were incubated with CHEX-seq probe (170 nM) in TES buffer (10 mM Tris, 1 mM EDTA, 150 mM NaCl) for 1 hr at room temperature. The samples were then washed with 1x PBS (w/o Ca^++^, Mg^++^) 3 times for 3 min.

#### Imaging and photoactivation

After CHEX-seq probe annealing and washing, the samples were transferred to the imaging chamber with 1x PBS (w/o Ca^++^, Mg^++^). All images and photoactivations were performed using a Carl Zeiss 710 Meta confocal microscope (20x water-immersion objectives, NA 1.0). CHEX-seq probe annealing was confirmed by exciting at 633 nm and emission was detected at 640-747 nm. The photoactivation was performed using the 405 nm (UV) laser at 60% power and 6.30 µs per pixel.

#### First strand DNA synthesis *in situ* and single cell harvest

After photoactivation in each individual cell’s nucleus, a master mix containing DNA polymerase I and 1^st^ strand DNA synthesis buffer was added to the cells and incubated for 1 h at room temperature. Subsequently, the single cells containing synthesized complementary DNA were harvested using a glass micropipette under using a Zeiss 710 confocal microscope (Carl Zeiss) for visualization.

#### Linear amplification of nucleosome free area of chromatin

(A) 1^st^ strand DNA synthesis and poly G tailing at 3’ end: After harvesting single cells, the *in situ* synthesized cDNA was removed by adding fresh prepared 0.1 N NaOH and incubating the sample for 5 min at RT followed by neutralization with 1 M Tris (pH 7.5). After ethanol precipitation, the 1st strand DNA was resuspended in nuclease free water. Subsequently, poly(G) was added to the 3’ end using terminal deoxynucleotidyl transferase (TdT) (Invitrogen). (B) 2^nd^ strand DNA synthesis and round 1 linear RNA amplification: 2^nd^ strand DNA was synthesized using DNA polymerase I for 2 h at 16 °C after priming with custom App-RC-polyC primer (Supplemental Data, Table 1). RNA was amplified using linear in vitro transcription from T7 RNA polymerase promoter incorporated into the double-stranded DNA with Ambion MEGAscript T7 In Vitro Transcription (IVT) Kit. (C) Round 2 1^st^ and 2^nd^ strand DNA synthesis and PCR amplification: After cleanup IVT reaction, 1^st^ strand DNA was reverse transcribed from aRNA using Superscript III using a custom App-RC primer (Supplemental Data, Table 1) 2nd strand DNA was synthesized using DNA Polymerase 1 with a custom 18bpPBC1 primer (Supplemental Data, Table 1). Subsequently, the double-stranded blunt ended DNA was amplified using custom primers 18bpPBC1 / App-RC (Supplemental Data, Table 1) following PCR condition: 98 °C for 30 sec; thermocycling at 98 °C for 10 sec, 50 °C for 30 sec, 72 °C for 30 sec for 27 cycles; extension at 72 °C for 2 mins, and was then used for library construction. Samples for the control experiments were processed with the same procedure except no CHEX-seq probe was applied, and 2^nd^ round 2^nd^ strand DNA PCR amplification was performed with custom primers 18bpPBC14 / App-RC (Supplemental Data, Table 1).

#### Sequencing Library Preparation

Illumina TruSeq Nano DNA Library Preparation Kit was used with modifications. All of the second round PCR amplified double-stranded DNA was used as input. After converting DNA fragment into blunt ends with End Repair Mix, base “A” was added; sequence adapters were ligated. DNA inserts were amplified with PCR.

#### External Data

*<u>GRO-seq</u>*: K562 GRO-seq was downloaded from SRA (accession GSE60454) (*42*) in FASTQ format; raw reads were processed using the SCAP-T pipeline (www.scap-t.org); POL2 engaged transcripts were inferred by HOMER (*43*); *ATAC-seq*: 1. Single-cell untreated K562 ATAC-seq data were downloaded from SRA (accession GSE65360) (*5*) in raw FASTQ format. We followed the alignment and peak calling methods in Buenrostro and colleagues (*5*); 2. Mouse brain ATAC-seq data were downloaded from ENCODE (Davis et al. 2018) (accession see Table S1) in BAM format; narrow and broad peaks were called using MACS2(*44*); *<u>DNase-seq</u>*: 1. K562 DNase-seq narrow and broad peaks were downloaded from ENCODE (accession see Table S1 SuppMaterials.pptx) in bigBed format; 2. Human brain DNase-seq data were downloaded from ENCODE (accession see Table S1) in BAM format; *<u>FAIRE-seq</u>*: K562 FAIRE-seq narrow peaks were downloaded from ENCODE (accession ENCFF000TLT) in BED format; the original hg19 genome build was lifted over to hg38 by CrossMap (*45*); *Reduced representation bisulfite sequencing* (*<u>RRBS)</u>*: K562 DNA methylation RRBS data were downloaded from UCSC ENCODE track in BEDMethyl format; the original hg19 genome build was lifted over to hg38 by CrossMap. *<u>ChIP-seq</u>*: K562 ChIP-seq data were downloaded from ENCODE (accession see Table S1) in genome build hg38. They were further organized in three categories: transcription factor binding sites (TFBSs) and narrow and broad histone modifications (H3K27ac, H3K4me3, H3K9ac, H3K4me2, H2AFZ; H3K4me1, H3K27me3, H3K36me3, H3K9me3, H3K79me2, H3K9me1). Only replicated peaks were used for histone modifications. *<u>Hi-C</u>*: K562 Hi-C data were downloaded from GSE63535 (*46*) in genome build hg19. In order to compare it with hg38 while minimizing potential artifacts caused by lifting over Hi-C data, we lifted over CHEX-seq from hg38 to hg19 using CrossMap. *<u>Enhancer and super-enhancer</u>*: Human and mouse experimentally validated enhancers were downloaded from the VISTA database (*47*); Super-enhancer data were downloaded from dbSUPER (*48*); *<u>DNA replication origin</u>*: K562 DNA replication origin data was downloaded from GEO (accession GSE46189), in BED format with pre-called peaks by the authors. The original genome build is hg19, which was converted to hg38 by CrossMap. *Enhancer/promoter interactions*: in UCSC Genome Browser, enhancers, promoters and regulatory interactions were loaded from database GeneHancer v4.11(*49*), using only high-confidence (“double elite”) data.

## Supplemental Data Tables

**Supplemental Data Table 1.**
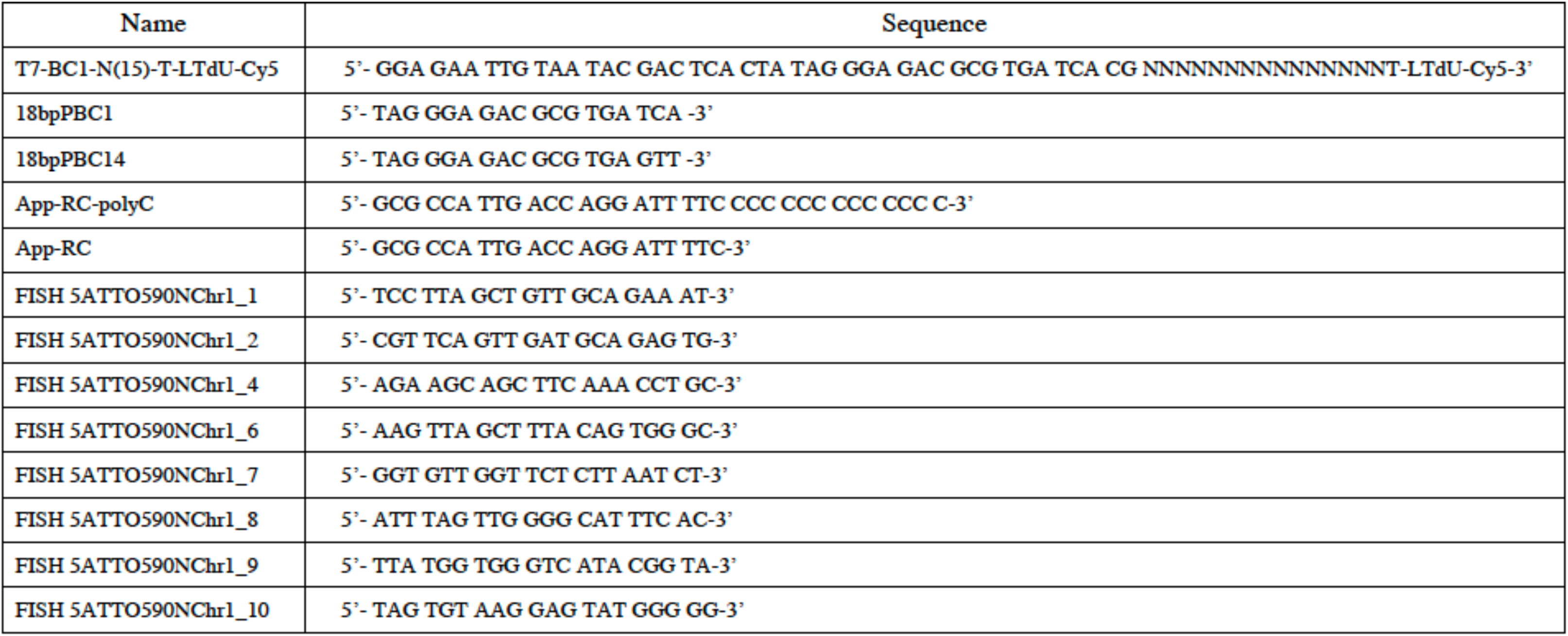
List of primers and oligonucleotide sequences used in these studies.

**Supplemental Data Table 2.**
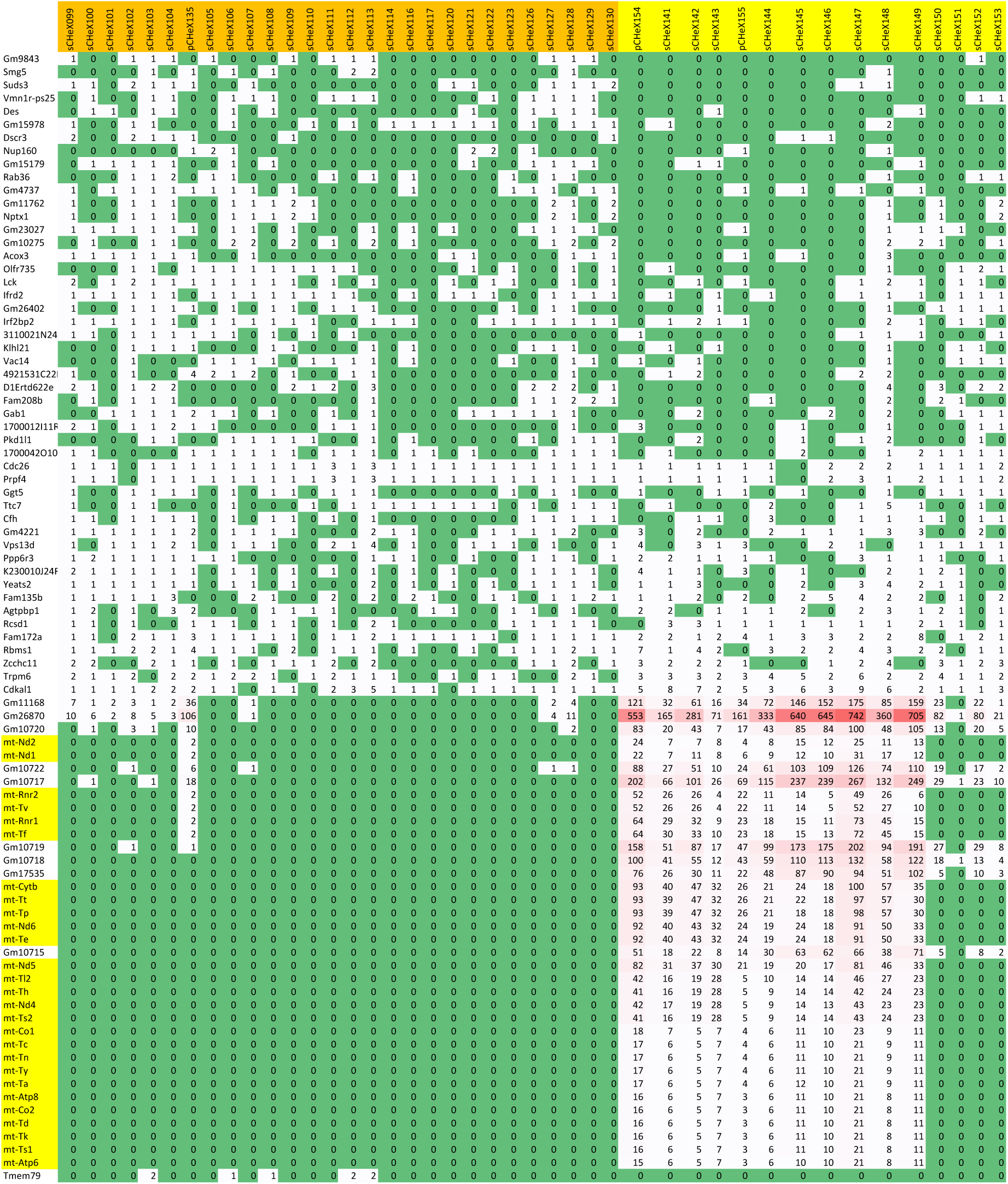
Genes that are differentially primed in single mouse neurons analyzed in tissue sections and dispersed cell culture. The proportional test was used to identify differentially CHEX-primer primed genes with BH corrected p-value of <0.05. For each gene, the group-wise sum is computed, and then compared with the grand sum where all genes are pooled. The Fisher exact test was applied to the contingency table to test the proportions.

## Supplemental Data Figures

**Supplemental Data Figure 1.**
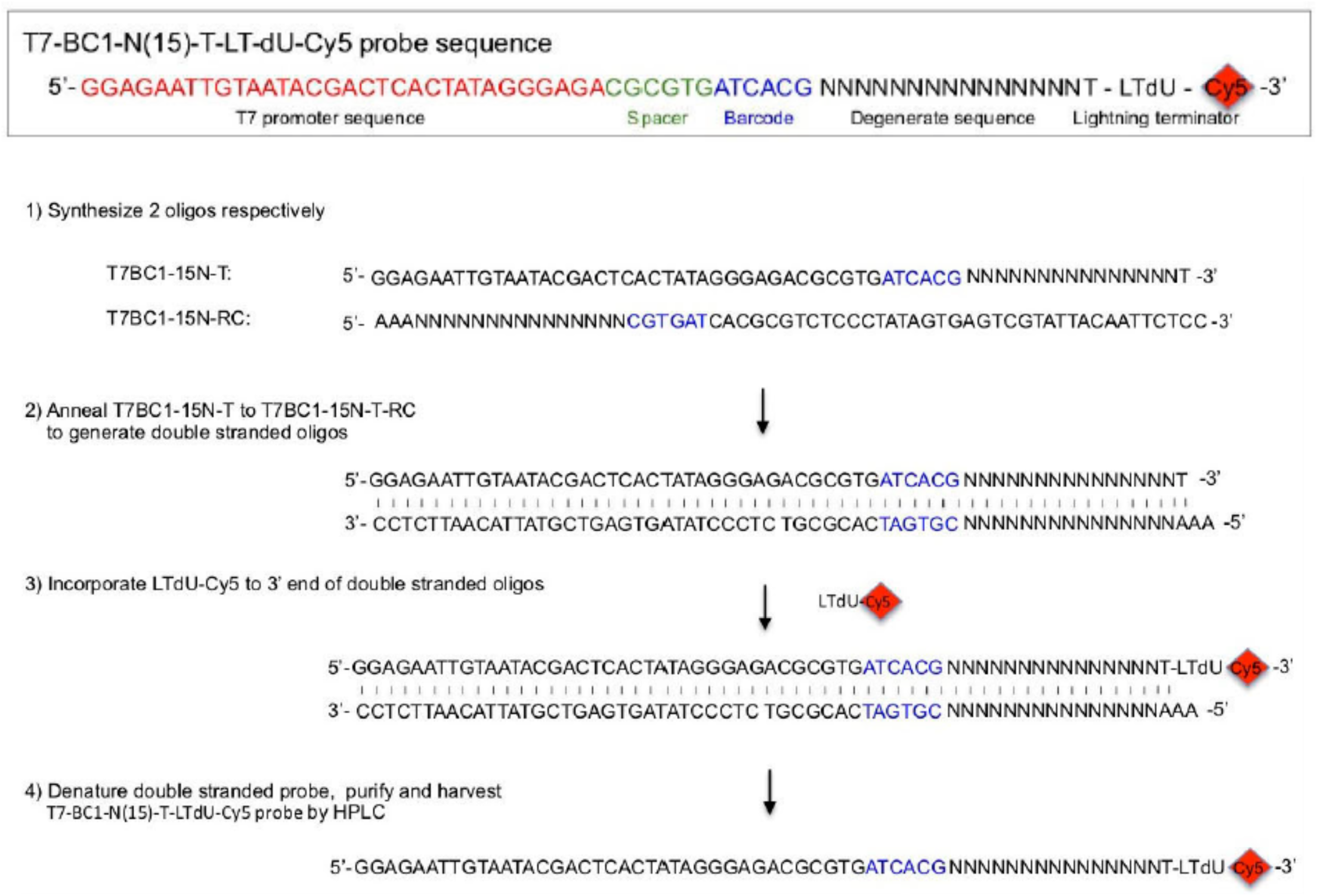
Structural scheme of CHEX-seq probe. The complete CHEX-seq probe (T7-BC1-N(15)-T-LTdU-Cy5) sequence is shown on the top. An oligo containing T7 promoter site, an Illumina 6 bp barcode1 (BC1, blue) and a 15 bp degenerate sequence, T7BC1-15N-T and its reverse complement oligo, T7BC1-15N-T-RC are synthesized and annealed to each other to generate double-stranded oligos. Cy5-labeled Lightning Terminator, LTdU-Cy5 is incorporated to its 3’ end. The single-stranded CHEX-seq probe is harvested after denaturation of the double-stranded probe and HPLC purification.

**Supplemental Data Figure 2.**
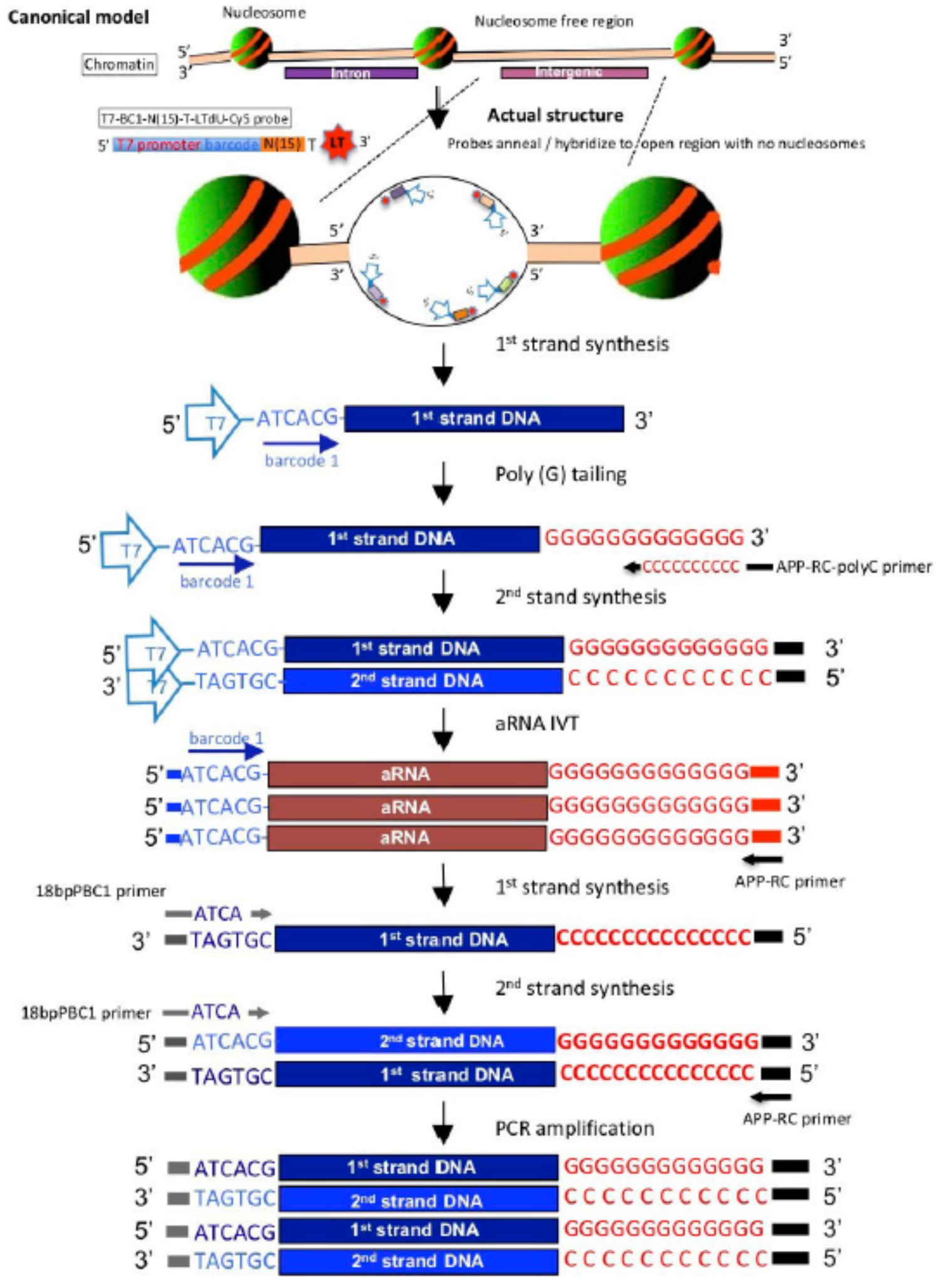
A schematic of the CHEX-seq aRNA amplification protocol. Upon applying the CHEX-seq probe, T7-BC1-N(15)-T-LTdU-Cy5, to the PFA fixed, Triton X-100 permeabolized cells, the degenerate N(15) sequence hybridizes to single-stranded nucleosome-depleted genomic DNA found within open chromatin regions. After laser-mediated photo-cleavage of the termination group of the CHEX oligonucleotide first strand DNA synthesis is primed by DNA polymerase I. Second strand DNA is primed and synthesized using custom App-RC-polyC primer (Supplemental Data, Table 1) after poly (G) tailing of 3’ end of 1^st^ strand DNA. Finally, RNA is amplified using linear in vitro transcription from the T7 RNA polymerase promoter incorporated into the double-stranded DNA. 2^nd^ round 1^st^ and 2^nd^ strand DNA subsequently are synthesized and amplified by PCR.

**Supplemental Data Figure 3.**
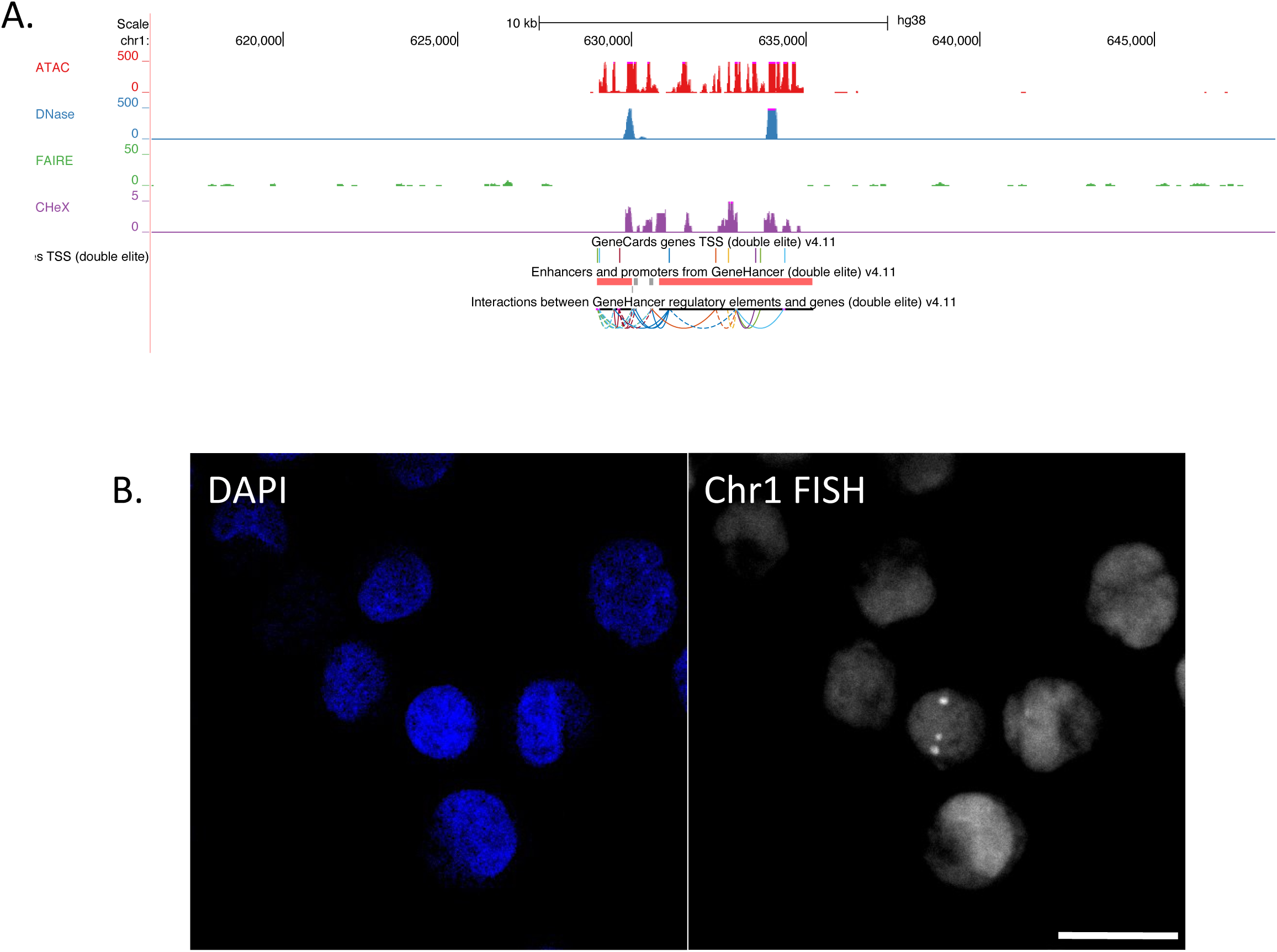
*In situ* hybridization to region 630737-633960 of chromosome 1 (hg38). The UCSC Genome Browser track view for a portion of chromosome 1 is shown in Panel A. The CHEX-seq track is similar to the ATAC-seq track showing that this chromosomal area is open. This is distinct from DNAse-seq and FAIRE-seq data. In Panel B, the left panel is the DAPI staining of the K562 cell nuclei. The right panel shows the fluorescence *in situ* hybridization signal using 8 fluorescently labeled oligonucleotides. These data show highly specific chromosome 1 trisomy in the K562 cells’ nuclei. Scale bar= 20µm.

**Supplemental Data Figure 4.**
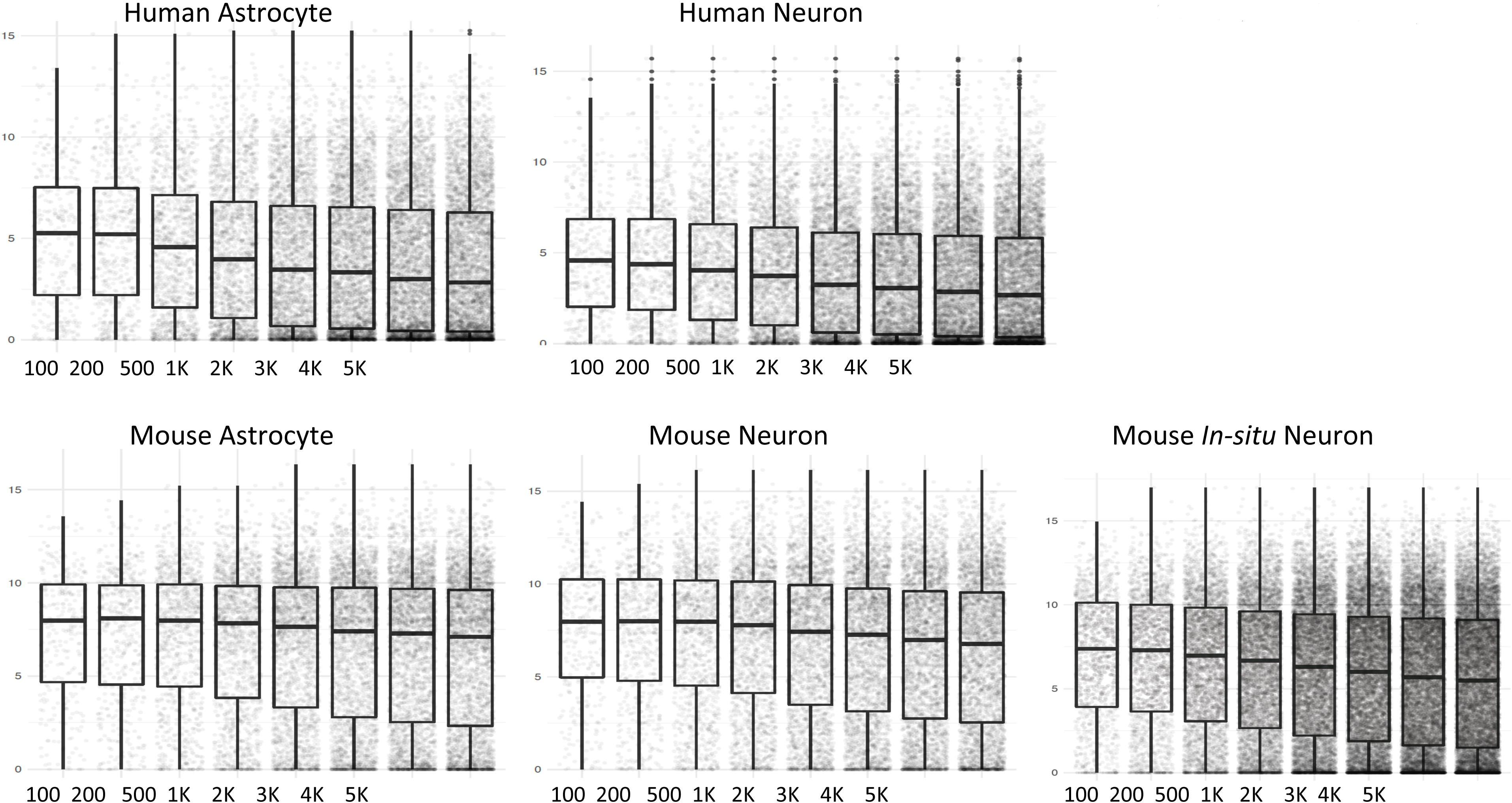
Correlation of CHEX-seq read distance from TSS with RNA abundance in neurons and astrocytes.

**Supplemental Data Figure 5.**
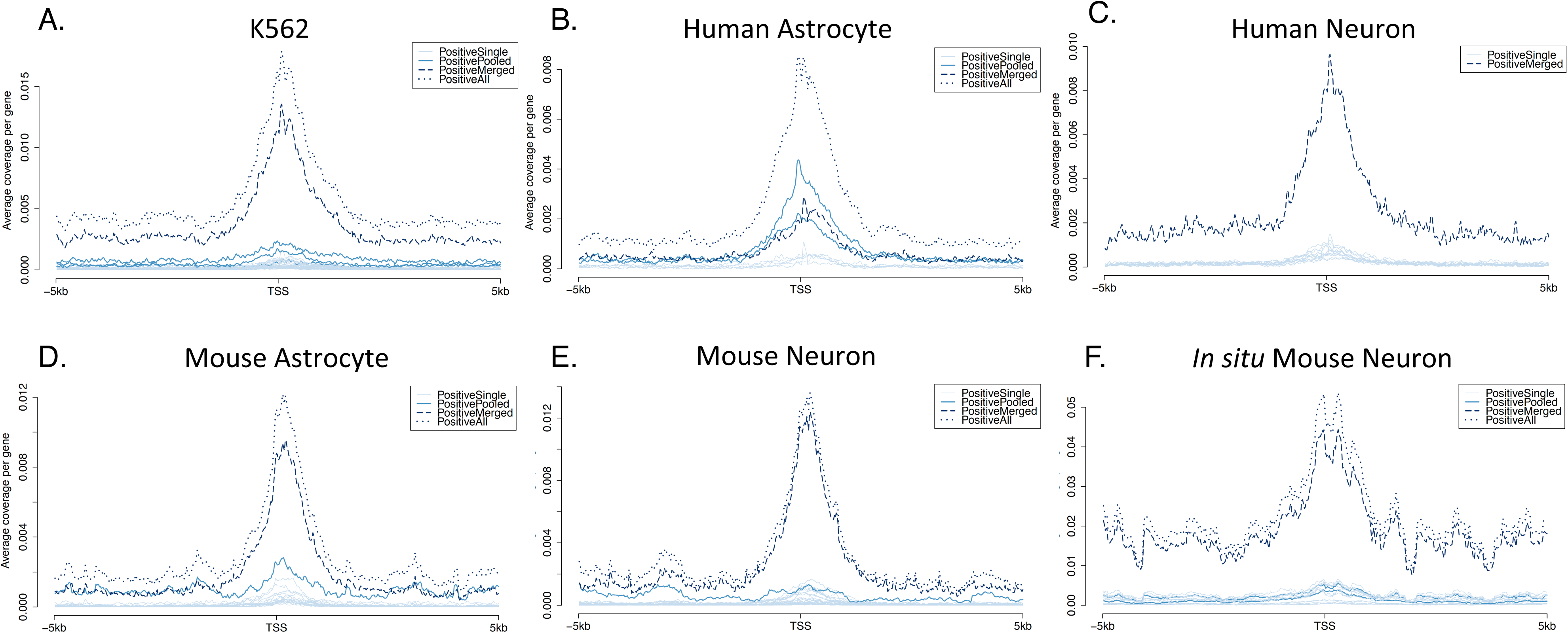
CHEX-seq reads mapping to the Transcriptional Start Sites for K562 cells, human and mouse dispersed neurons and astrocytes and mouse brain section localized neurons.

**Supplemental Data Figure 6.**
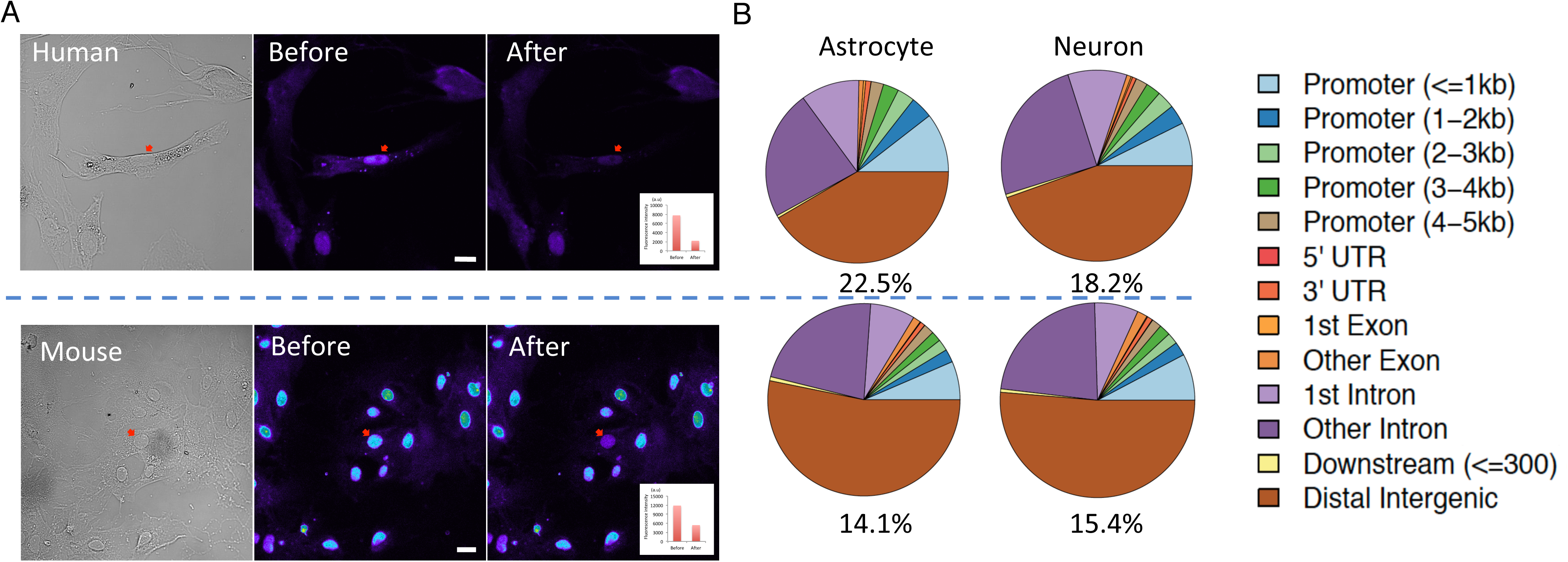
CHEX-seq applied to primary astrocyte cultures from mouse and human samples. (A) Images of human astrocytes on top and mouse on bottom. DIC (left) and DAPI images (left and middle panels, respectively) before CHEX-seq probe activation, and DAPI image after activation (right panels; quantification of DAPI signal in insert). scale bar= 20µm. (B) Quantification of CHEX-seq priming sites with respect to genomic features in astrocytes (left) and neurons (right). Key for CHEX-seq read site of localization relative to gene structure.

